# Longitudinal diffusion tensor imaging identifies coordinated white matter remodeling during cocaine self-administration in mice

**DOI:** 10.64898/2026.10.14.759212

**Authors:** Zachary Fernandez, Alice McQueney, Melinda Osowski, Xiang Liu, Yun Liang, Shuangtao Ma, Andrew Bender, Norman Scheel, Erik Shapiro, Alfred J. Robison, Chunqi Qian

## Abstract

White matter microstructural alterations are increasingly recognized as a feature of cocaine use disorder (CUD), yet the longitudinal progression and biological underpinnings of these changes remain poorly understood. Although rodent intravenous self-administration (IVSA) provides a translational model of voluntary cocaine use, longitudinal diffusion tensor imaging (DTI) has not previously been combined with cocaine IVSA in mice. Here, we established a longitudinal IVSA-DTI framework to characterize white matter microstructural remodeling during cocaine self-administration and early withdrawal. Male C57BL/6 mice underwent DTI at baseline, following 11 consecutive days of cocaine (0.5 mg/kg/infusion) or saline IVSA, and after a 7-day withdrawal period immediately following a cocaine-seeking test (n = 4/group). Fractional anisotropy (FA) was evaluated longitudinally using atlas-based region-of-interest analyses and whole-brain diffusion maps. Cocaine-treated mice acquired robust operant responding and exhibited coordinated longitudinal FA alterations across interconnected hippocampal-associated white matter pathways. The most consistent changes occurred within the bilateral dorsal hippocampal commissure, fornix, and fimbria, with cocaine-treated mice demonstrating increased FA during self-administration relative to baseline while saline controls exhibited smaller or opposing longitudinal changes. Difference-in-differences maps further demonstrated cocaine-associated FA alterations within hippocampal pathways and localized regions of the corpus callosum after accounting for longitudinal changes in saline controls. These findings establish the feasibility of combining longitudinal DTI with cocaine IVSA in mice and demonstrate coordinated white matter microstructural remodeling during voluntary cocaine exposure. This framework provides a translational platform for future studies integrating longitudinal neuroimaging with genetically modified mouse models, tractography, and histological validation to investigate the mechanisms underlying cocaine-associated white matter plasticity.

## Introduction

Addiction, which affects an estimated 48.5 million Americans, is associated with structural brain changes that are reinforced through repeated substance use. Diffusion tensor imaging (DTI) provides a noninvasive approach to assess white matter microstructure by measuring the diffusion of water molecules within neural tissue, enabling investigation of the structural circuitry underlying addiction. Clinical DTI studies and preclinical animal models have consistently demonstrated white matter alterations following cocaine exposure and dependence (Lim et al., 2008; Tondo et al., 2021). For example, individuals with cocaine use disorder (CUD) exhibit altered white matter integrity within the genu and rostral body of the corpus callosum (CC) (Moeller et al., 2005). Furthermore, longitudinal studies have shown that altered fractional anisotropy (FA), a sensitive DTI marker of white matter microstructure, is associated with greater cocaine use and poorer clinical outcomes (Ma et al., 2017; Xu et al., 2010). These findings link cocaine-associated alterations in brain structure with drug-seeking behavior, relapse vulnerability, and long-term functional impairment.

Preclinical animal models have similarly identified cocaine-associated white matter alterations. For instance, experimenter-administered cocaine decreases FA within the splenium of the CC in male rats (Narayana et al., 2009). However, because experimenters control drug dose, frequency, and duration in most paradigms, they do not fully capture the voluntary nature of human cocaine use. In contrast, intravenous self-administration (IVSA) allows rodents to voluntarily obtain cocaine through operant responding and is widely considered the gold-standard preclinical model of CUD. Importantly, when combined with DTI, IVSA in rats revealed alterations within both the genu and splenium of the CC that more closely resemble findings observed in human cocaine users than those reported following experimenter-administered cocaine (Gozzi et al., 2010; Ma et al., 2009; Moeller et al., 2005; Narayana et al., 2009).

Despite these advances, longitudinal DTI has not been combined with cocaine IVSA in mice. Establishing this approach is important because mice provide substantial experimental advantages over rats, including lower cost, higher throughput, and access to numerous transgenic lines that permit mechanistic interrogation of specific genes, cell types, and neural circuits (Jaramillo et al., 2014). Recent advances in jugular catheter design, small-animal magnetic resonance imaging (MRI) acquisition, and diffusion image processing have now made longitudinal IVSA-DTI studies in mice technically feasible (Deloulme et al., 2024; Micheau et al., 2025; Valles et al., 2022). Nevertheless, no study has combined longitudinal DTI with IVSA to characterize white matter changes across baseline, cocaine self-administration, and withdrawal in the same animals. Moreover, previous DTI studies of cocaine IVSA have been performed almost exclusively in rats, leaving the longitudinal effects of cocaine self-administration on mouse white matter largely uncharacterized.

In the present study, we established a longitudinal mouse IVSA-DTI framework by acquiring diffusion MRI before cocaine exposure, following 11 days of cocaine self-administration, and after a 7-day withdrawal period immediately following a cocaine-seeking test. Longitudinal changes in FA and MD were evaluated within white matter pathways previously implicated in addiction while also generating whole-brain diffusion maps for regional visualization. In this small cohort, the most prominent and consistent longitudinal alterations were observed in FA; therefore, the present report focuses on FA-based analyses of cocaine-associated white matter remodeling. This study establishes the feasibility of longitudinal IVSA-DTI in mice, provides the first characterization of cocaine-associated white matter remodeling in a mouse IVSA model, and establishes a foundation for future mechanistic investigations into the cellular and molecular basis of these structural adaptations.

## Methods

### Animals

Male C57BL/6 mice, twelve weeks of age at the beginning of the experiment, underwent longitudinal DTI at three time points: before IVSA (baseline), after 11 consecutive days of cocaine or saline self-administration, and after a seven-day withdrawal period. The final withdrawal imaging session was performed one day after a cocaine-seeking test (**Figure 1**). The present study included four cocaine-treated and four saline-treated mice. All cocaine and saline mice completed 11 total days of daily self-administration sessions, but due to a premature death only three cocaine and four saline mice completed the seeking test. All eight mice completed baseline and post-self-administration imaging. Following the premature death of one cocaine-treated mouse, seven animals completed the behavioral seeking test. The deceased animal was subsequently imaged post-mortem. Mice were housed at 22–25 °C on a 12 h light–dark cycle with food and water ad libitum, except for where noted. Behavioral testing took place during the light phase. All experiments were approved by Michigan State University Institutional Animal Care and Use Committee and were carried out in accordance with the guidelines set in the Guide for the Care and Use of Laboratory Animals of the National Institutes of Health.

**Figure 1.**
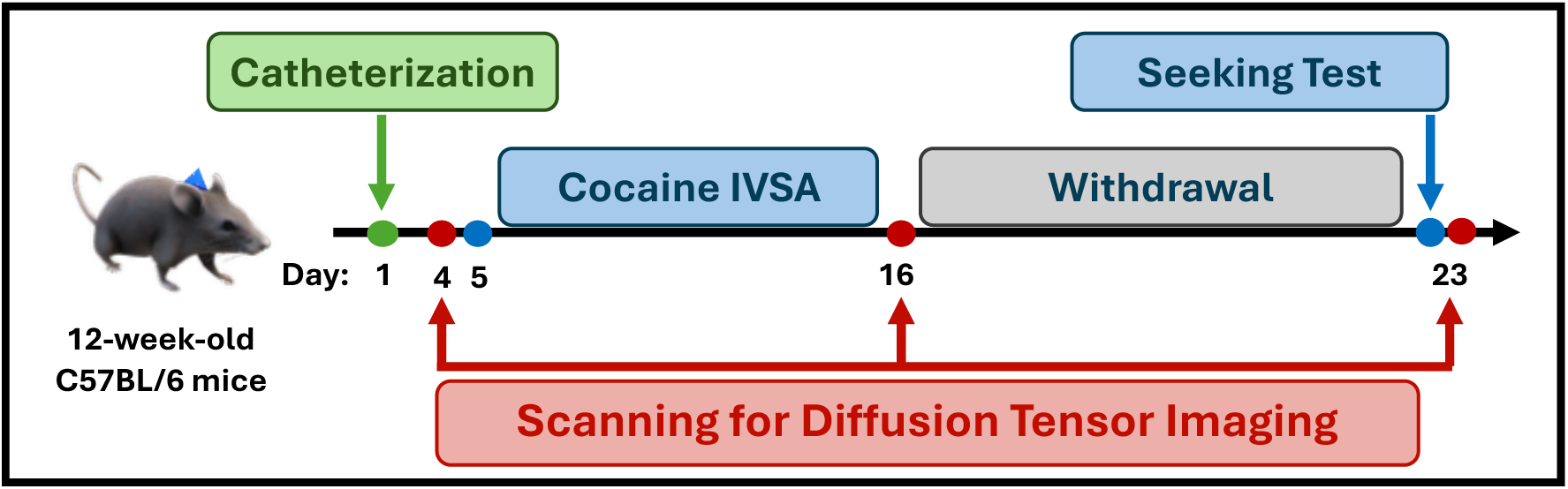
*Study timeline.* Experimental design illustrating diffusion tensor imaging (DTI) acquisition at baseline, following 11 days of intravenous cocaine self-administration (IVSA), and after a 7-day withdrawal period immediately following a cocaine-seeking test. Mice underwent jugular vein catheterization prior to behavioral training.

### Jugular catheterization

Mice underwent surgical implantation of a chronic indwelling jugular vein catheter as previously described (Doyle et al., 2021; Valles et al., 2022). Mice were anesthetized with isoflurane, and a catheter (Instech) was implanted into the right jugular vein and secured with two ligatures. The catheter attached subcutaneously to an MRI-compatible vascular access button (Instech) that sat between the animal’s shoulder blades. This allowed for repeated intravenous cocaine or saline delivery without interfering with MRI acquisition during the scan sessions (**Figure 2A**). Animals were allowed four days to recover before beginning self-administration. Prior to testing and after the last day of daily self-administration, patency was assessed by infusion of 0.02mL of Brevital Sodium (5mg/mL, i.v.). Catheters were deemed patent if a rapid loss of muscle tone and/or a ‘frozen’ stance was observed within 3-5 seconds of infusion.

**Figure 2.**
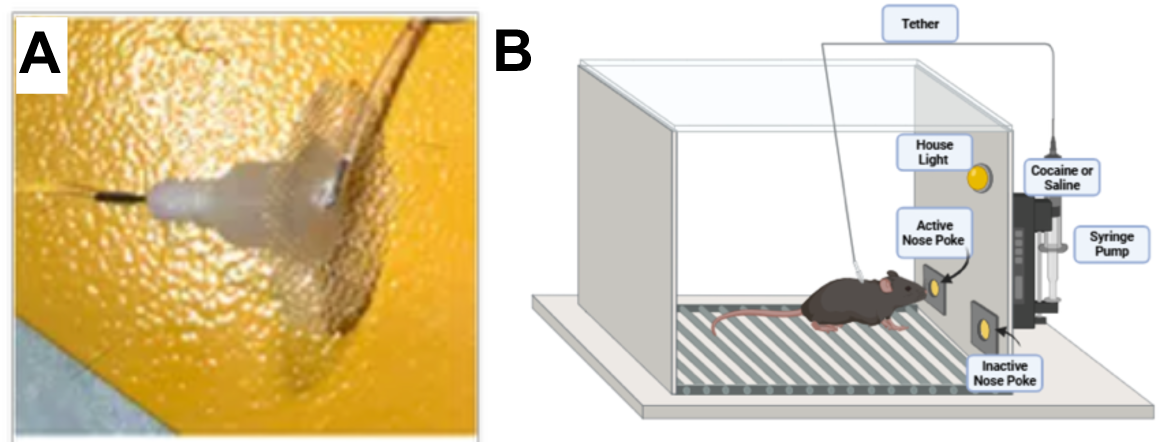
*Experimental paradigm for cocaine IVSA.* (A) MRI-compatible vascular access button used for repeated intravenous drug delivery. (B) Operant conditioning chamber used for cocaine self-administration.

### Cocaine IVSA

Beginning four days after catheterization, mice underwent daily two-hour IVSA sessions for 11 consecutive days on a fixed ratio one schedule of reinforcement. Operant chambers (Med-Associates Inc.) contained two illuminated nose-poke ports designated as active or inactive, counterbalanced across subjects (**Figure 2B**). Mice were randomly assigned to respond for cocaine or saline, and each group had an equal amount of left versus right active nose pokes. Mice were food-deprived overnight before the first self-administration session and maintained at 90% free-feeding weight until the last day of daily self-administration. The active nose-poke was baited with sucrose pellets during the first two sessions.

Responses in the active port resulted in activation of the syringe pump for a 1.6-second intravenous infusion of cocaine hydrochloride at 0.5 mg/kg per infusion, or an equivalent volume of saline. Each infusion was paired with the house light and active nose poke light turning off for 20 seconds to signal a timeout before turning back on. Responses in the inactive port were recorded but had no programmed consequence. Sessions were limited to a maximum of 75 infusions.

After the final self-administration session, mice remained undisturbed in their home cages for seven days. They were then returned to the operant chamber for a 1-hour cocaine-seeking test conducted under conditions matching the self-administration context. Active nose pokes produced the conditioned pump-associated cue but no infusion. Behavioral outcomes included active nose pokes, inactive nose pokes, and mock infusions earned during self-administration.

Daily self-administration behaviors (active nose pokes, rewards, inactive nose pokes) were analyzed with a two-way repeated measures ANOVA. Seeking test parameters (active nose pokes, mock rewards, and inactive nose pokes) were analyzed with a Welch’s t test.

### MRI acquisition

Diffusion MRI was performed longitudinally at baseline, after completion of the 11-day self-administration period and immediately following the cocaine-seeking test conducted after seven days of withdrawal (**Figure 1**). All images were acquired using a 7T Bruker BioSpin small-animal MRI system. A surface coil was positioned immediately anterior to the vascular access button and secured in place. Mice were induced with 5% isoflurane and maintained at 1–2% isoflurane in oxygen during image acquisition. Animals were positioned in a dedicated mouse imaging cradle, and respiration, heart rate, and body temperature were continuously monitored. Respiratory rate was maintained between approximately 50 and 70 breaths per minute.

Diffusion-weighted images were acquired using a spin-echo echo-planar imaging sequence with a repetition time of 2 seconds, echo time of 24.1 milliseconds, field of view of 19.2 × 19.2 × 16 mm, and voxel dimensions of 0.15 × 0.15 × 0.50 mm³. Diffusion weighting was applied at a b-value of 2047.72 s/mm² along 20 noncollinear diffusion-encoding directions. A high-resolution T2-weighted anatomical image was also obtained for anatomical registration.

### Image preprocessing

Diffusion-weighted images were processed using a combination of MRtrix3, FSL, AFNI, and in-house MATLAB scripts. Raw diffusion images underwent standard preprocessing, including Gibbs ringing removal, eddy current correction, and tensor fitting to generate voxel-wise maps of FA and mean diffusivity (MD). For each animal, the non-diffusion-weighted (b0) image was first registered to its corresponding high-resolution anatomical T2-weighted image using affine registration followed by nonlinear normalization. The anatomical image was subsequently normalized to the C57BL/6 mouse template, and the resulting transformations were applied to the diffusion-derived parameter maps. Atlas-based segmentation was performed using the DukeRCCF mouse brain atlas, allowing extraction of regional diffusion metrics from anatomically defined regions of interest (**Figure 3**). Mean FA and MD values were extracted from each ROI for every animal at each imaging time point prior to group-level analyses.

**Figure 3.**
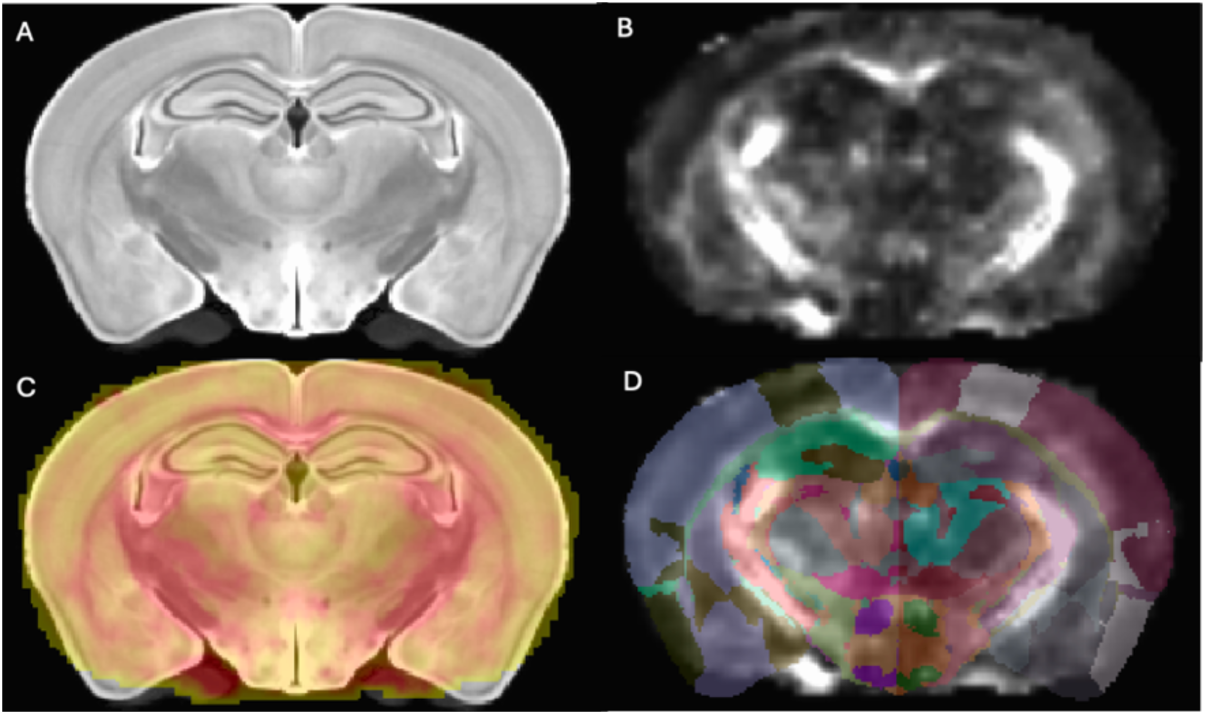
*DTI processing and atlas registration pipeline.* (A) Representative T2-weighted anatomical image. (B) Corresponding fractional anisotropy (FA) map generated from DTI. (C) Registered anatomical image with FA overlay demonstrating image alignment. (D) Atlas-based segmentation overlaid on the registered FA image for extraction of region-specific diffusion metrics.

### Region-of-Interest and Subtraction Map Analysis

Although diffusion metrics were extracted throughout the brain, the present study focused primarily on white matter pathways because the interpretation of FA and MD is most established within highly organized fiber tracts. Longitudinal FA and MD values were initially evaluated within white matter pathways previously implicated in cocaine exposure and learning, including the DHC, fimbria, fornix, CC, and fasciculus retroflexus. Averaged diffusion metrics were extracted from each ROI at baseline, following 11 days of cocaine self-administration, and after the 7-day withdrawal period. FA demonstrated the most prominent and consistent longitudinal alterations in the present cohort and was therefore selected as the primary diffusion metric for the analyses and figures reported here.

To visualize longitudinal changes within each treatment group, subtraction images were generated by subtracting baseline FA maps from post-self-administration FA maps: ΔFA = FA _SA_ – FA _BL_. Positive values indicate increased FA relative to baseline, whereas negative values indicate decreased FA. Equivalent subtraction maps were generated for the withdrawal time point. To isolate cocaine-associated changes while accounting for longitudinal alterations observed in saline controls, a difference-in-differences calculation was performed: ΔFA _Cocaine_ – ΔFA _Saline_. In which, ΔFA represents the change in FA from baseline to self-administration. These cocaine-effect maps visualize longitudinal FA alterations observed in cocaine-treated mice relative to saline controls after accounting for longitudinal changes associated with repeated imaging and behavioral training (**Figure 8**).

### Comparison Between Groups

Behavioral data, including active nose pokes, inactive nose pokes, and rewards earned, are presented as mean ± standard error (SE). Regional diffusion metrics were calculated as mean ± SE for each experimental group at baseline (BL), self-administration (SA), and withdrawal (WD). Because FA demonstrated the most prominent and consistent longitudinal alterations in this preliminary cohort, the quantitative values and longitudinal trajectories presented in the current report focus on FA. For each ROI, longitudinal change scores were calculated between imaging sessions (ΔSA−BL, ΔWD−BL, and ΔWD−SA) to characterize the magnitude and direction of white matter microstructural remodeling. Group mean trajectories were visualized to examine temporal changes in interconnected white matter pathways.

Behavioral and diffusion measurements were analyzed to characterize longitudinal changes across imaging sessions. Given the preliminary cohort size, statistical analyses were exploratory, with emphasis placed on effect magnitude, longitudinal consistency across individual animals, and concordance between quantitative ROI analyses and representative subtraction maps.

## Results

### Cocaine self-administration produces robust acquisition of operant responding

Mice readily acquired cocaine IVSA over the 11-day training period (**Figure 4**). Cocaine-treated mice exhibited progressively greater active nose-poke responding and earned substantially more drug infusions than saline controls throughout training. There was a main effect of Subject, indicating that some mice had more active nosepokes (p= 0.0028, F(6, 60) = 3.81) and earned more rewards (p<0.0001, F(6,60) = 10.61) than others. The main effect of drug was not significant in nosepokes (p=0.1414) or rewards (p=0.1714), though this is likely due to the small sample size. In contrast, inactive nose pokes remained comparatively low in both groups, demonstrating that operant responding was directed toward the reinforced active port rather than generalized increases in exploratory behavior.

**Figure 4.**
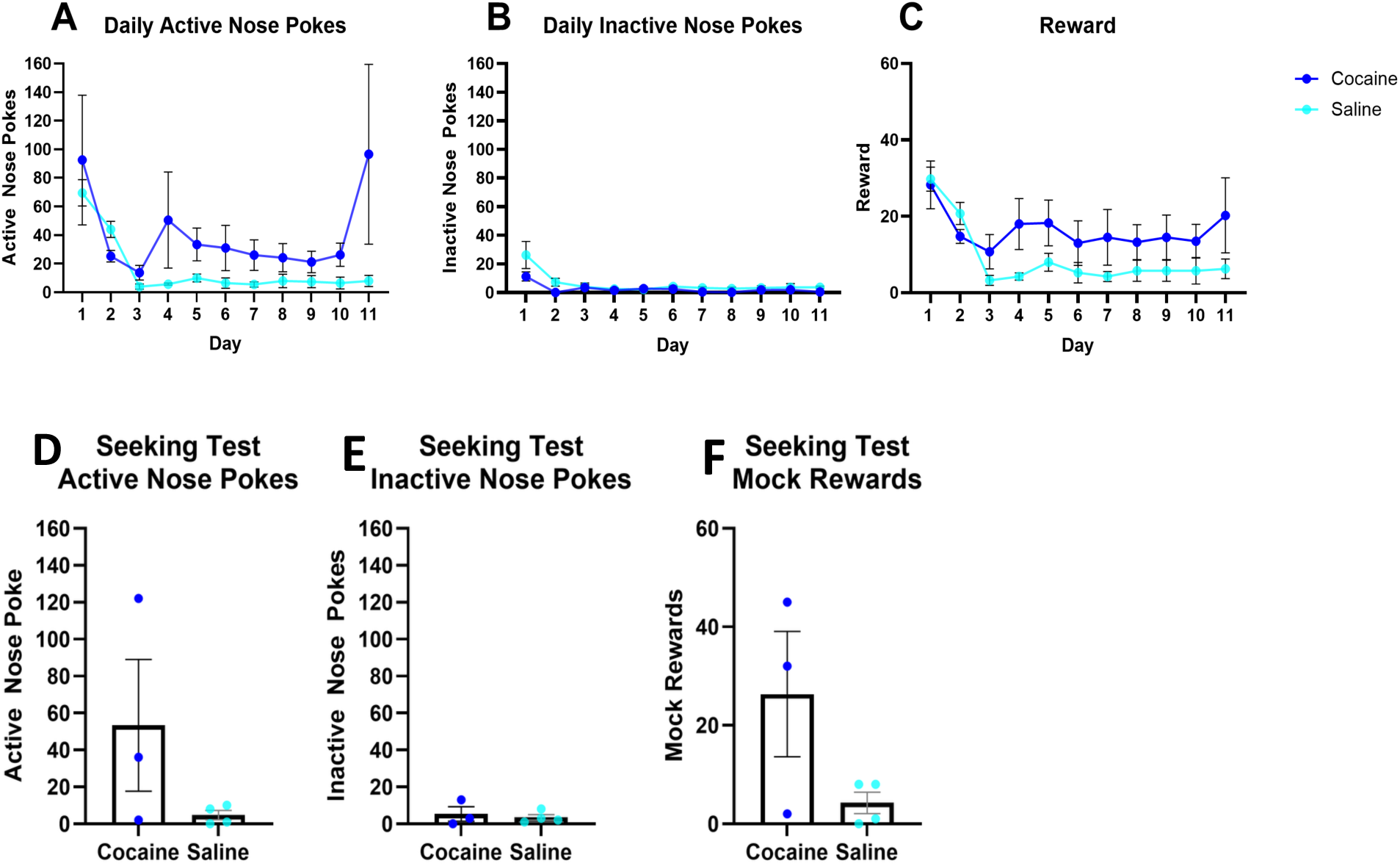
*Behavioral performance during IVSA and subsequent seeking test.* Longitudinal behavioral measures across training sessions showing (A) active nose pokes, (B), inactive nose pokes and (C) rewards (drug infusions) earned for cocaine and saline groups. Cocaine mice acquired operant responding, demonstrating increased active nose-poke and reward acquisition relative to saline controls. Following seven days of withdrawal, cocaine-seeking behavior was assessed by measuring (D) active nose pokes, (E) inactive nose pokes, and (F) mock rewards earned during a 1-hour seeking test conducted without cocaine delivery. Cocaine-treated mice exhibited greater average active nose-poke responding and mock reward acquisition than saline controls, although these differences did not reach statistical significance. Data are presented as mean ± standard error (SE).

In the seeking test (**Figure 4 D-F**), mice exposed to cocaine had a much higher average number of nosepokes and earned mock rewards, though these effects did not reach significance due to variance and small number of subjects. Together, these findings confirm successful acquisition of cocaine-reinforced operant behavior and establish the behavioral framework for subsequent longitudinal neuroimaging analyses.

### Longitudinal white matter remodeling following cocaine self-administration

FA and MD were initially evaluated to characterize longitudinal diffusion changes following cocaine self-administration. In this preliminary cohort, MD did not demonstrate longitudinal alterations of comparable magnitude or consistency, whereas FA demonstrated coordinated changes across multiple white matter pathways and was therefore the focus of subsequent analyses (**Figures 5 and 7**). The most consistent changes occurred within interconnected hippocampal-associated white matter tracts, including the DHC, fornix, and fimbria, whereas more modest changes were observed within the CC and fasciculus retroflexus.

**Figure 5.**
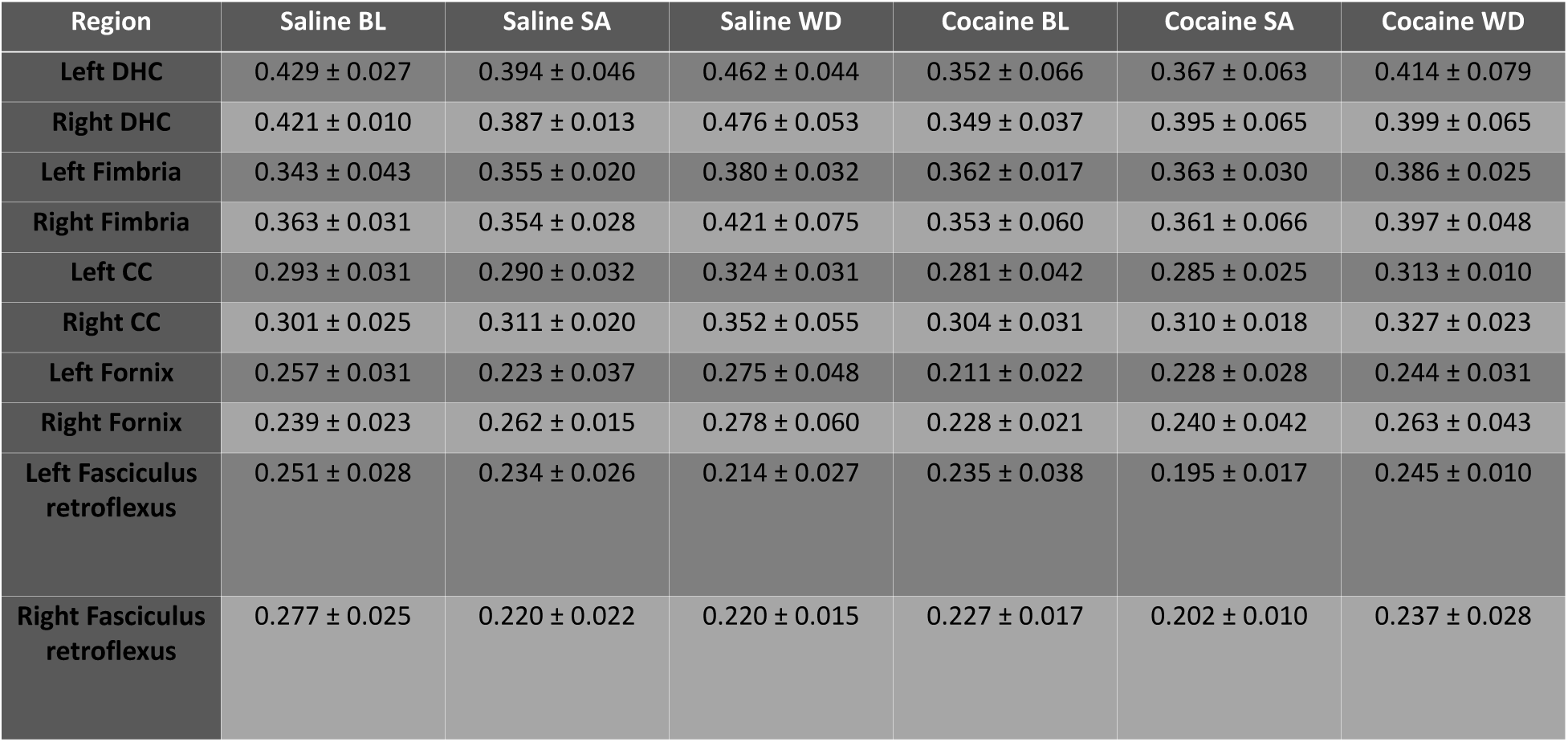
*Regional FA values across longitudinal imaging time points.* Mean ± SE FA values for selected white matter regions of interest in saline- and cocaine-treated mice at baseline (BL), following self-administration (SA), and after withdrawal (WD). Representative white matter regions include the dorsal hippocampal commissure (DHC), fimbria, corpus callosum (CC), fornix, and fasciculus retroflexus. The DHC, fimbria, and fornix represent interconnected hippocampal-associated white matter pathways involved in learning and memory, whereas the corpus callosum and fasciculus retroflexus were included because of their reported involvement in cocaine-associated structural remodeling.

**Figure 6.**
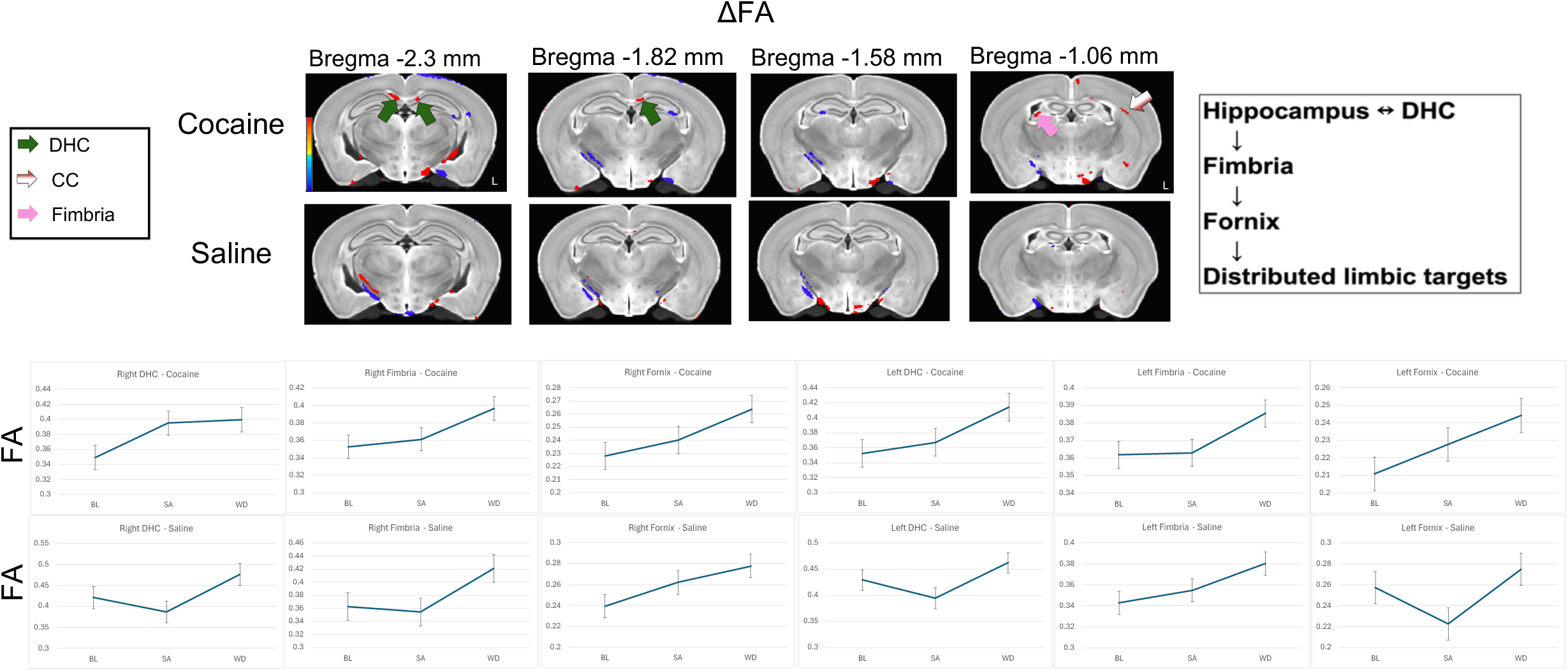
*Longitudinal white matter microstructural changes following cocaine self-administration*. Representative subtraction maps (ΔFA = FA_SA_ – FA_BL_) for cocaine and saline groups highlighting white matter regions demonstrating longitudinal FA alterations. Representative subtraction maps highlight the dorsal hippocampal commissure (DHC), fimbria, and corpus callosum (CC). Coronal sections are shown at approximate stereotaxic levels corresponding to the Paxinos and Franklin mouse brain atlas. Corresponding longitudinal ROI traces illustrate mean FA values across baseline, self-administration, and withdrawal, demonstrating coordinated increases in FA within hippocampal-associated white matter pathways following cocaine exposure.

**Figure 7.**
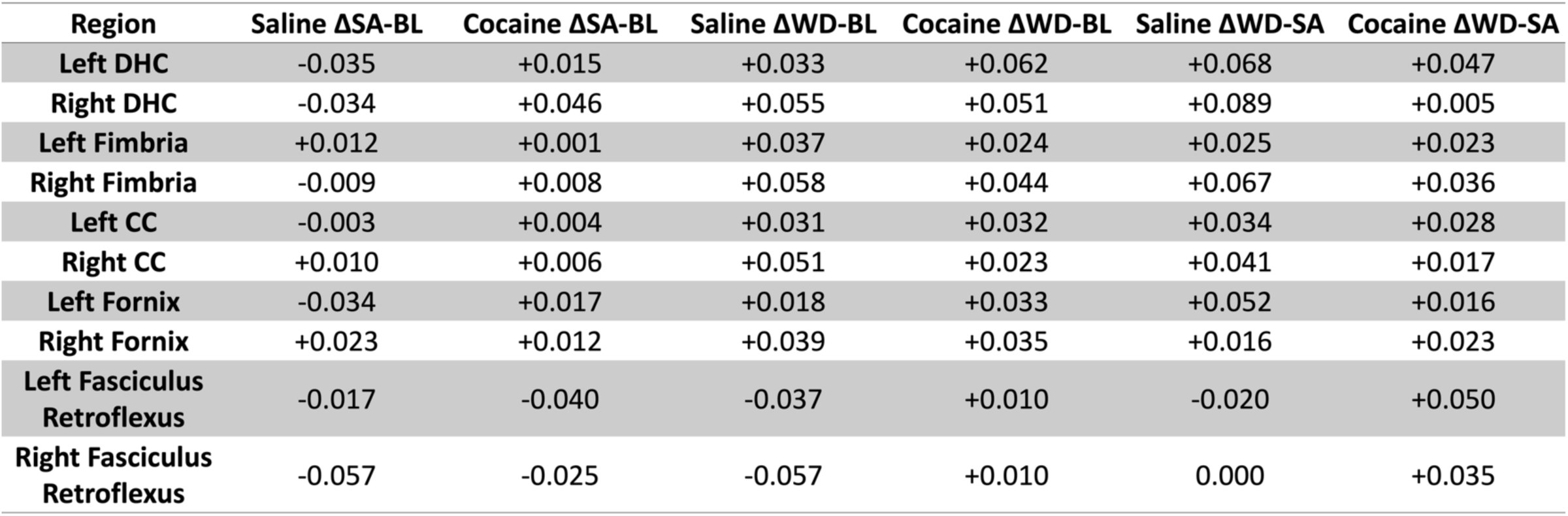
*Magnitude of longitudinal FA changes between imaging time points.* Mean longitudinal changes in fractional anisotropy (ΔFA) for selected white matter regions between baseline and self-administration (ΔSA–BL), baseline and withdrawal (ΔWD–BL), and withdrawal relative to self-administration (ΔWD–SA) for saline and cocaine groups. Positive values indicate increased FA between imaging sessions, while negative values indicate decreased FA. Regions include the dorsal hippocampal commissure (DHC), fimbria, fornix, corpus callosum (CC), and fasciculus retroflexus. The hippocampal-associated pathways (DHC, fimbria, and fornix) demonstrated the most consistent longitudinal alterations following cocaine self-administration.

**Figure 8.**
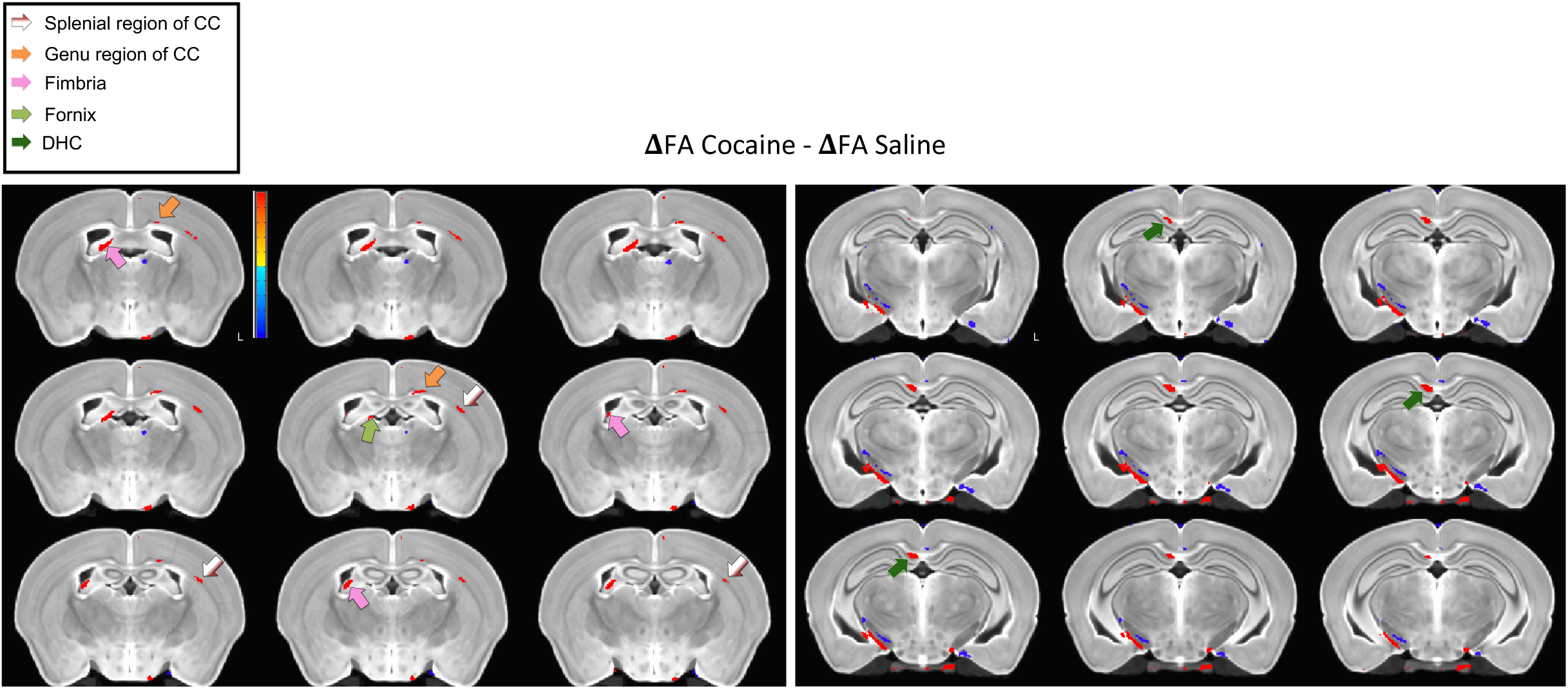
*Cocaine-specific white matter remodeling following self-administration.* Difference-in-differences subtraction maps illustrating the cocaine effect on fractional anisotropy: ΔFA _Cocaine_ – ΔFA _Saline_, in which ΔFA = FA _SA_ – FA _BL_. Representative slices highlight cocaine-associated alterations within the dorsal hippocampal commissure (DHC), fimbria, and genu and splenium of the corpus callosum (CC), relative to saline controls.

During the self-administration phase, cocaine-treated mice generally exhibited increased FA relative to baseline across these hippocampal pathways, while saline-treated animals demonstrated comparatively smaller changes or decreases in FA. The bilateral DHC demonstrated the clearest longitudinal separation between treatment groups, with cocaine-treated mice exhibiting increased FA during self-administration whereas saline controls generally exhibited reductions relative to baseline. Similar longitudinal trajectories were observed within the bilateral fornix and fimbria, forming a coordinated pattern across anatomically connected hippocampal-associated white matter pathways. These observations were corroborated by the quantitative ROI measurements (**Figure 5**), which demonstrated consistent increases in mean FA across these pathways in cocaine-treated animals.

Many of these alterations remained evident at the withdrawal time point, although saline-treated mice also exhibited longitudinal changes during withdrawal, reducing the separation between groups compared with the self-administration time point. Consequently, the most robust cocaine-associated differences were observed immediately following self-administration, suggesting that these changes may be transient and driven by acute drug effects.

### Longitudinal white matter trajectories support coordinated hippocampal pathway remodeling

Longitudinal subtraction maps generated from the baseline and self-administration time points (ΔFA = FA _SA_ – FA _BL_) demonstrated localized FA increases within several white matter pathways following cocaine self-administration (**Figure 6**). Prominent changes were observed within the DHC, fimbria, and localized regions of the CC.

Subject-level ROI trajectories demonstrated consistent longitudinal increases in FA across the bilateral DHC, bilateral fornix, and bilateral fimbria in cocaine-treated mice (**Figure 6**). In contrast, saline controls exhibited more heterogeneous trajectories, with several pathways demonstrating transient decreases during self-administration followed by partial recovery during withdrawal. The consistent directional changes observed across anatomically connected hippocampal white matter pathways demonstrate a coordinated network-level pattern of microstructural remodeling associated with cocaine self-administration rather than isolated regional effects.

### Difference-in-differences analysis isolates cocaine-associated white matter alterations

To distinguish cocaine-associated changes from longitudinal alterations observed in saline controls, difference-in-differences subtraction maps were generated by subtracting saline longitudinal FA changes from those observed in cocaine-treated mice (**Figure 8**). These cocaine-effect maps highlighted localized FA increases within the DHC, fimbria, and anterior and posterior regions of the CC after accounting for changes associated with repeated imaging and behavioral training. The spatial distribution of these cocaine-effect maps closely paralleled the quantitative ROI analyses, providing complementary evidence of greater cocaine-associated FA alterations within interconnected hippocampal-associated white matter pathways in this preliminary cohort. Together, the behavioral analyses, quantitative ROI measurements, longitudinal trajectory analyses, and cocaine-effect subtraction maps consistently demonstrate coordinated remodeling of interconnected hippocampal-associated white matter pathways following cocaine self-administration.

## Discussion

The present study establishes the feasibility of combining longitudinal DTI with IVSA in mice to characterize white matter microstructural remodeling across operant drug exposure and withdrawal. This system could be generalized to observe changes in white matter occurring at different doses of cocaine, using progressive ratio responding, comparing sexes, or changes in response to self-administration of other drugs, including opiates. Critically, this system demonstrates feasibility of the use of mice rather than rats in this context, bringing to bear all of the genetic tools available in mice to perform mechanistic and causal studies of the effects of cocaine on white matter organization.

Cocaine-treated mice successfully acquired operant responding, demonstrating increased active nose pokes and cocaine infusions relative to saline controls. Longitudinal DTI revealed coordinated alterations in FA within interconnected hippocampal-associated white matter pathways, including the bilateral DHC, fornix, and fimbria. These findings were consistently supported by ROI analyses, longitudinal trajectory plots, and representative subtraction maps, collectively demonstrating a spatially coherent pattern of cocaine-associated white matter remodeling.

Previous human neuroimaging studies have consistently identified white matter abnormalities among individuals with CUD, including frontal and callosal alterations detected using DTI (Lim et al., 2008; Ma et al., 2009, 2017; Moeller et al., 2005; Tondo et al., 2021; Xu et al., 2010). Likewise, both experimenter-administered cocaine and IVSA studies in rats have demonstrated alterations within the genu and splenium of the CC (Gozzi et al., 2010; Narayana et al., 2009). The present findings extend this literature by demonstrating that longitudinal DTI can be successfully combined with cocaine IVSA in mice. Although localized changes within the CC were observed, the most consistent alterations occurred within hippocampal-associated white matter pathways, suggesting that cocaine-associated structural remodeling extends beyond the CC and involves distributed limbic circuitry engaged during cocaine reinforcement. This interpretation is consistent with functional neuroimaging studies showing disrupted mesocorticolimbic and frontostriatal connectivity in individuals with CUD (Gu et al., 2010; Wilcox et al., 2011). The present findings therefore extend previous rat IVSA studies by identifying hippocampal-associated white matter pathways as additional regions exhibiting longitudinal microstructural remodeling during cocaine self-administration.

One of the most notable findings was the coordinated increase in FA across multiple anatomically connected hippocampal-associated white matter pathways during cocaine self-administration. The DHC, fornix, and fimbria collectively form major interhemispheric and output pathways of the hippocampus, a structure that plays a central role in associative learning, contextual memory, and cue-reward processing (Fuchs et al., 2005). During IVSA, mice learned to discriminate between active and inactive nose-poke ports, with active responses producing cocaine delivery paired with conditioned sensory cues. The hippocampus has been implicated in contextual encoding and retrieval of cocaine-associated memories, making these pathways relevant for the acquisition and expression of drug-seeking behavior (Eagle et al., 2026; Fuchs et al., 2005; See, 2005). More broadly, altered connectivity within distributed memory systems has been associated with differences in memory performance across neuroimaging studies, supporting the importance of considering coordinated network-level changes rather than regional effects (Viviano et al., 2019). In contrast, saline-treated animals exhibited substantially lower levels of operant responding because saline lacked the reinforcing properties of cocaine. The coordinated longitudinal changes observed across these pathways therefore may reflect experience-dependent structural remodeling associated with the acquisition of cocaine-reinforced drug-seeking behavior rather than nonspecific effects of repeated cocaine exposure alone (Fields, 2015; Scholz et al., 2009), potentially involving neural circuits supporting drug-related learning and memory and contributing to subsequent drug-seeking behavior during abstinence. Together, these findings suggest that cocaine-associated white matter remodeling preferentially involves pathways supporting learned associations and contextual memory rather than uniform changes throughout the brain.

Although increased FA is often interpreted as reflecting improved white matter organization, diffusion tensor imaging cannot distinguish among the multiple biological processes that influence water diffusion (Jones et al., 2013). Alterations in FA may reflect changes in myelination, axonal density, fiber coherence, extracellular space, or neuroinflammatory processes rather than a single underlying cellular mechanism. Consequently, FA should be interpreted as a sensitive, but nonspecific, marker of white matter microstructure rather than a direct measure of myelin integrity (Jones et al., 2013).

Accordingly, the present findings should be interpreted as evidence of cocaine-associated white matter microstructural remodeling rather than direct evidence of altered myelination. Future histological studies determining whether myelination is altered in these regions after cocaine IVSA are needed.

MD was also evaluated in the present cohort but did not demonstrate comparable longitudinal alterations to those observed in FA. The more prominent FA alterations observed in the present dataset may reflect differential sensitivity of these diffusion metrics to the underlying microstructural changes; however, the modest sample size precludes conclusions regarding metric-specific biological effects. Evaluation of additional cohorts will determine whether additional MD alterations emerge with increased statistical power.

## Limitations

Several limitations should be considered when interpreting these findings. First, this study represents a preliminary longitudinal cohort consisting of four cocaine-treated and four saline-treated mice. Accordingly, the analyses were exploratory and interpreted primarily according to effect magnitude, longitudinal consistency across individual animals, and agreement between quantitative ROI analyses and representative subtraction maps. Nevertheless, the convergence of multiple complementary analyses supports the robustness of the observed longitudinal patterns. The modest sample size may also limit sensitivity to smaller longitudinal effects in other diffusion metrics, including MD, which did not demonstrate changes of comparable magnitude in the present cohort. Second, atlas-based segmentation was performed using the DukeRCCF mouse brain atlas because it provides robust anatomical registration for diffusion MRI analyses. However, its CC segmentation is limited to left and right hemispheric regions rather than anatomically distinct genu, body, and splenium subdivisions. Consequently, localized CC alterations previously reported in rat IVSA studies may have been underrepresented in the present analyses (Gozzi et al., 2010; Narayana et al., 2009). Future studies incorporating higher-resolution white matter parcellations may provide greater anatomical specificity for evaluating regional CC remodeling. Third, while DTI provides sensitive measures of white matter microstructure, diffusion metrics are not biologically specific and require histological validation to identify the underlying cellular mechanisms responsible for the observed imaging changes. Finally, tractography and whole-brain structural connectome analyses are ongoing and will further characterize cocaine-associated network remodeling in future studies. Future studies combining diffusion MRI with histological markers of myelin, microglia, and astrocytes will be important for identifying the cellular substrates underlying these imaging findings.

## Conclusions

In sum, this study establishes the feasibility of longitudinal DTI in a mouse IVSA model and demonstrates coordinated remodeling of interconnected hippocampal-associated white matter pathways during cocaine exposure. These findings establish a translational platform for future studies integrating longitudinal DTI with genetically modified mouse models to investigate the mechanisms underlying cocaine-associated white matter remodeling.

## Acknowledgements

This study was supported by the American Heart Association (DIVSUP1077212) to Z.F., by NIH/NINDS (RF1NS128611) to C.Q, and by NIH/NIDA (R01DA040621) to A.J.R. The authors would like to thank Missi Rogers for her assistance on surgical procedures as well as Michael Henin, Bleon Jahiri, Anja Lee, Nivi Medum, Charvi Bagewadi Ellur, Sophie Peret, Ishitha Teluguntla and Maxwell Wickerson for contributions to MR image preprocessing for this study.

## Abbreviations

DTI: diffusion tensor imaging
MRI: magnetic resonance imaging
CUD: cocaine use disorder
IVSA: intravenous self-administration
FA: fractional anisotropy
MD: mean diffusivity
ROI: region of interest
DHC: dorsal hippocampal commissure
CC: corpus callosum
BL: baseline
SA: self-administration
WD: withdrawal
SE: standard error
EPI: echo-planar imaging
TE: time of echo
TR: time of repetition
FOV: field of view
FSL: FMRIB Software Library
AFNI: Analysis of Functional NeuroImages

